# Inhibition of the TRIM24 bromodomain reactivates latent HIV-1

**DOI:** 10.1101/2022.06.16.496524

**Authors:** Riley M. Horvath, Zabrina L. Brumme, Ivan Sadowski

## Abstract

Expression of the HIV-1 genome by RNA Polymerase II is regulated at multiple steps, as are most cellular genes, including recruitment of general transcription factors and control of transcriptional elongation from the core promoter. We recently discovered that tripartite motif protein TRIM24 is recruited to the HIV-1 Long Terminal Repeat (LTR) by interaction with TFII-I and causes transcriptional elongation by stimulating association of PTEF-b/ CDK9. Because TRIM24 is required for stimulation of transcription from the HIV-1 LTR, we were surprised to find that IACS-9571, a specific inhibitor of the TRIM24 C-terminal bromodomain, induces HIV-1 provirus expression in otherwise untreated cells. IACS-9571 reactivates HIV-1 in T cell lines bearing multiple different provirus models of HIV-1 latency. Additionally, treatment with this TRIM24 bromodomain inhibitor encourages productive HIV-1 expression in newly infected cells and inhibits formation of immediate latent transcriptionally repressed provirus. IACS-9571 synergizes with PMA, ionomycin, TNF-α and PEP005 to activate HIV-1 expression. Furthermore, co-treatment of CD4^+^ T cells from individuals with HIV-1 on antiretroviral therapy (ART) with PEP005 and IACS-9571 caused robust provirus expression. Notably, IACS-9571 did not cause global activation of T cells; rather, it inhibited induction of IL2 and CD69 expression in human PBMCs and Jurkat T cells treated with PEP005 or PMA. These observations indicate the TRIM24 bromodomain inhibitor IACS-9571 represents a novel HIV-1 latency reversing agent (LRA), and unlike other compounds with this activity, causes partial suppression of T cell activation while inducing expression of latent provirus.

## Introduction

The global HIV pandemic has persisted for 40 years [1]. Despite intensive research over this time, an estimated 38 million people are currently living with HIV-1, the vast majority of whom will require anti-retroviral therapy (ART) for the remainder of their lives [2]. Current anti-retroviral therapy (ART) for HIV-1 can suppress virus replication to undetectable levels, but does not affect the population of latently infected cells that harbor transcriptionally silenced provirus that develop during the course of infection [3]. These latently infected cells represent a major barrier for development of a cure for HIV-1 infection, and consequently a variety of therapeutic strategies to eliminate these cells are currently under intense investigation [4] [5].

One potential strategy that has garnered considerable attention, initially designated “Shock and Kill” would involve intervention to force reactivation of latent provirus such that latently infected cells become exposed to the host immune response and/ or additional therapeutic means for their elimination [6]. This potential strategy has prompted development and characterization of small molecule drugs and compounds that are capable of inducing expression of latent provirus, collectively referred to as latency reversing agents (LRAs). LRAs characterized to date inhibit a variety of functions that impact regulation of transcription from the HIV-1 LTR, including histone modification, transcriptional elongation, and signal transduction [7].

The 5’ long terminal repeat (LTR) of integrated HIV-1 provirus contains numerous *cis*-elements for host cell factors that respond to multiple cellular signaling pathways involved in immune cell stimulation as well as growth factors and cytokines [8]. Considering the number and variety of transcription factors capable of binding the HIV-1 LTR, many of which have transcriptional activation function [9], it is remarkable that this virus invariably develops transcriptionally silenced provirus in resting memory T cells and monocyte-derived macrophages. Latent provirus is produced by multiple pathways in unstimulated cells, involving layers of regulation including, but not limited to down regulation of transcription factors responsive to immune signaling pathways, binding of transcriptional repressors to the 5’ LTR that recruit histone deacetylases (HDACs) and histone methyltransferases (HMTs), and subsequent recruitment of silencing complexes that promote spreading of repressive chromatin marks from the silenced LTR [10]. Consequently, HIV-1 expression in cells that revert to a resting state becomes shut down over a period of days through a combination of epigenetic silencing and additional mechanisms that include loss of the viral transactivator protein TAT. Additionally, ∼50% of cells newly infected with HIV-1 produce a population of provirus that is transcriptionally repressed immediately (Dahabieh *et al*, 2013). This mode for production of latency was denoted early or immediate latency, and appears to involve novel mechanism(s) that are influenced by signaling downstream of the T-cell receptor (Dahabieh *et al*. 2014) and possess the capability of bypassing the function of TAT (Hashemi *et al*, 2016), and typified by association of YY1 with the 5’ LTR [11].

Latent HIV-1 provirus in resting memory T helper cells becomes reactivated in response to T-cell receptor (TCR) engagement with antigen presenting dendritic cells. TCR signaling stimulates the activity of multiple transcriptional activators capable of binding the 5’ HIV-1 LTR which cause recruitment of general transcription factors necessary for transcription of the provirus genome [12]. Like many cellular genes, the 5’ LTR promoter of latent HIV-1 is associated with RNA Pol II that is paused shortly after initiation because of inhibition by the factors NELF and DSIF [13] [14]. Consequently, a key event for reactivation of HIV-1 provirus involves recruitment of viral TAT protein which binds nascent HIV-1 TAR RNA associated with paused RNA Polymerase II. TAT directly recruits the P-TEFb complex containing CDK9, which phosphorylates the pausing factors NELF and DSIF, and RNA Pol II CTD S2 to promote transcriptional elongation [15]. Reactivation of HIV-1 expression in response to TCR signaling requires multiple factors specifically bound to the 5’ LTR, including TFII-I which is constitutively associated with two highly conserved cis-elements flanking the LTR enhancer region in association with USF1/2 [16] [17]. We discovered that TFII-I recruits the coactivator protein Tripartite-Motif containing protein 24 (TRIM24) to the LTR, and this interaction promotes transcriptional elongation from the LTR core promoter by RNA Polymerase II [18], an effect that is associated with enhanced recruitment of CDK9 and increased RNA Pol II CTD on serine 2 phosphorylation. These observations indicate that TRIM24 may function as a transcriptional coactivator by promoting recruitment of P-TEFb.

TRIM24, previously designated TIF-1α, was initially discovered as a co-factor for transcriptional activation by nuclear hormone receptors [19]. Like other TRIM protein family members, TRIM24 contains conserved N-terminal RING, B-Box zinc finger, and coiled-coil domains, in addition to C-terminal PHD (plant homeodomain) and bromodomain motifs, which bind the histone H3 with the combination of unmodified K4 and acetylated K23 epigenetic marks [20]. Overexpression of TRIM24 is associated with poor prognosis of a variety of cancers including breast, non-small cell lung, hepatocellular and glioblastomas [21][22], but overall the molecular mechanism(s) by which this factor regulates transcription has not been elucidated. Specific inhibitors that target bromodomains have been developed, as this motif is considered to be “druggable” [23]. Using protein structure guided design, a TRIM24 bromodomain binding compound, IACS-9571, was developed with nanomolar affinity that inhibits interaction with histone H3K23Ac peptides [24]. Interestingly, IACS-9571 was found to inhibit cell growth and metastatic invasive potential of glioblastomas from patients [22]. These observations support the contention that TRIM24 may contribute to cancer progression through interaction of chromatin through its bromodomain.

In this study we examine the effect of IACS-9571 on HIV-1 transcription, and surprisingly, find this compound causes reactivation of latent provirus in multiple different T cell line models of provirus latency, and produces synergistic effects in combination with T cell signaling agonists. Treatment of T cells with IACS-9571 inhibits production of immediate latent provirus, and encourages productive expression of HIV-1 in newly infected cells. IACS-9571 treatment caused elevated association of TRIM24 with the LTR suggesting that inhibition of the bromodomain may cause redistribution of this factor from cellular genes. Treatment of CD4^+^ T cells from individuals with HIV-1 on ART with a combination of PEP005 and IACS-9572 resulted in robust viral transcription, while also suppressing expression of T cell activation markers IL2 and CD69. These results indicate that the TRIM24 bromodomain inhibitor IACS-9571 represents a novel class of latency reversing agent which may prove useful for strategies to purge latently infected cells.

## Results

### The TRIM24 bromodomain inhibitor IACS-9571 promotes HIV-1 expression

We previously discovered that TRIM24 is recruited to the HIV-1 LTR by interaction with TFII-I, and this causes enhanced elongation from the HIV-1 LTR through elevated CDK9 recruitment and RNA Pol II CTD S2 phosphorylation [18]. Importantly, Jurkat cell lines bearing *TRIM24* gene knockouts are defective for reactivation of latent HIV-1 in response to T cell signaling. Consequently, we examined what effect the TRIM24 bromodomain inhibitor might have on expression of HIV-1, using a cell line which bears an HIV-1 mini-virus reporter where luciferase is expressed as a fusion with Gag (Fig. 1A). Surprisingly, rather than inhibiting HIV-1 expression, IACS-9571 treatment caused activation of HIV-1 luciferase expression in a dose-dependent manner, where treatment with 25 μM caused significant 3-fold induction in otherwise untreated cells (Fig. 1B). In parallel experiments we found that this compound produced minimal toxicity at concentrations where we observed significant reactivation (Fig. 1C). A similar effect of IACS-9571 was observed using the JLat10.6 cell line, which bears a full-length HIV-1 provirus where *Nef* has been replaced by GFP (Fig. S1).

**Figure 1.**
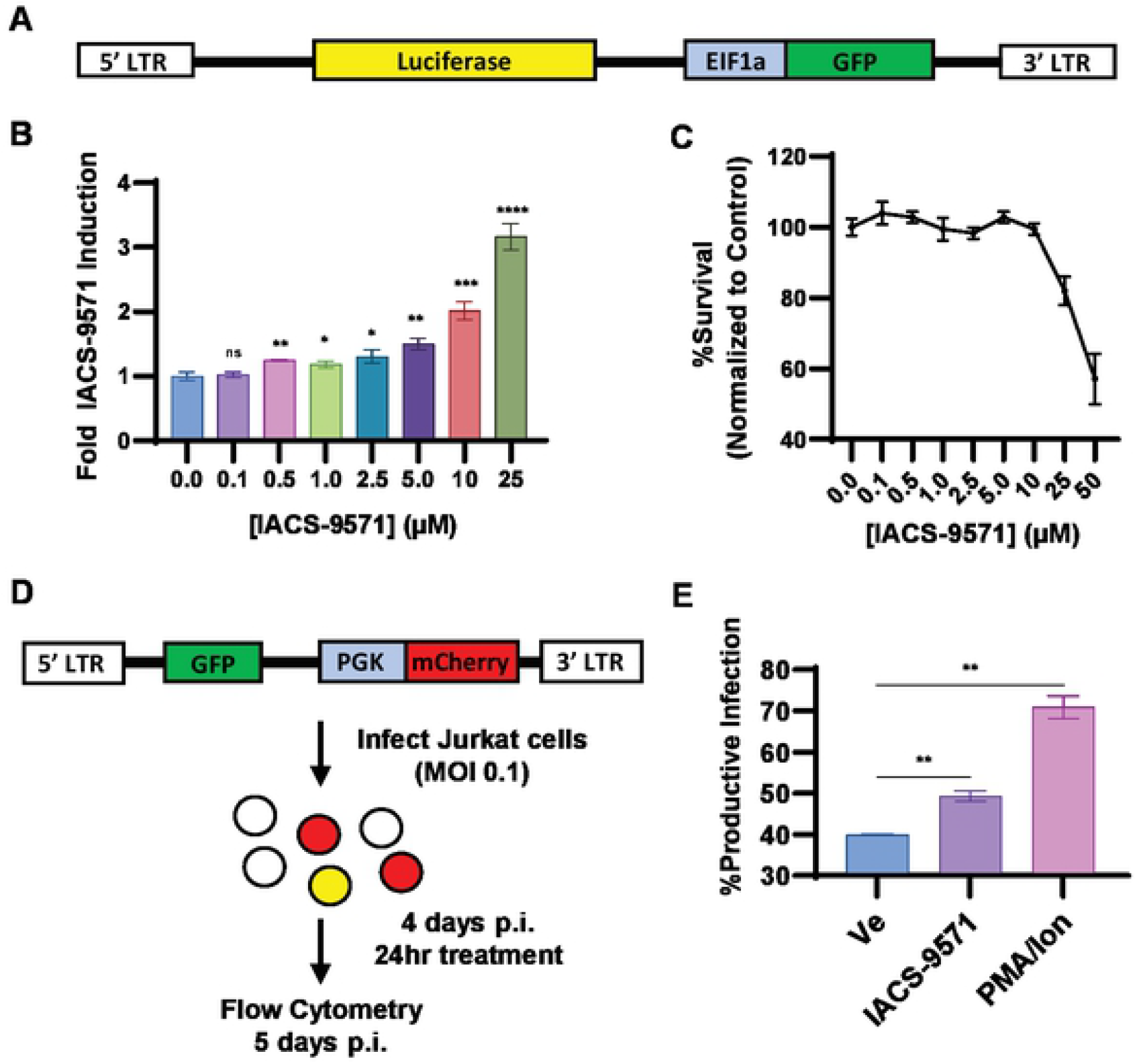
IACS-9571 promotes HIV-1 expression. **Panel A:** Schematic representation of HIV-1 mini-virus in the KB1 Jurkat Tat cell line, where Luciferase is expressed from the 5’ LTR as a fusion with Gag. **Panel B:** KB1 cells were treated with the indicated concentration of IACS-9571 for 4 hrs prior to luciferase assay. Results are an average of three determinations, and error bars represent standard deviation. **Panel C:** KB1 cell viability was determined after 4 hrs treatment with IACS-9571 at the indicated concentration. Viability assays were performed in duplicate with error bars representing standard deviation shown. **Panel D:** Depiction of Red-Green-HIV-1 (RGH) reactivation assays. RGH is a full-length replication incompetent HIV-1 virus with GFP expressed by the 5’ LTR as a fusion with Gag and *Nef* is replaced with a PGK promoter that drives mCherry expression [42]. Wildtype Jurkat T cells were infected with RGH at a low MOI; 4 days post-infection (p.i.), pools of cells were treated with 10 μM IACS-9571, 20 nM PMA/ 1 μM ionomycin, or left untreated for 24 hrs. Subsequently, productive and latently infected cells were analyzed by flow cytometry. **Panel E:** Ratio of productively infected cells following RGH reactivation. Results are an average of two determinations, and error bars represent standard deviation.

We also examined the effect of IACS-9571 shortly following HIV-1 infection using a dual reporter Red-Green HIV (RGH) which enables detection of infected cells by expression of mCherry from an internal PGK promoter, and independent measurement of expression from the 5’ LTR with a GFP reporter (Fig. 1D). With this reporter virus, productively infected cells express both mCherry and GFP, whereas infected cells that produce immediate latent provirus only express mCherry (Fig. 1*D*). Treatment of RGH infected cells with a combination of PMA and ionomycin 4 days post-infection produced ∼70% productive infections, a significant increase from the ∼40% productive infections in DMSO (Ve) treated cells (Fig. 1*E*). Interestingly, we also observe a similar increase in the proportion of productively infected cells, albeit smaller, caused by IACS-9571 compared to the untreated control (Fig. 1*E*), indicating that this TRIM24 bromodomain inhibitor is capable of causing activation of latent provirus in a significant proportion of newly infected T cells.

Additionally, we examined the effect of IACS-9571 on establishment of latency or productive replication upon initial infection. Untreated Jurkat T cells or cells treated with 10 μM IACS-9571, were infected with the RGH reporter virus and the proportion of productively infected cells, indicated by expression of both GFP and mCherry was monitored for 10 days post-infection (Fig. 2*A*). Consistent with previous observations, we found that ∼35% of infected untreated cells (Ve) developed productive infections at 4 days post infection, as indicated by expression of GFP; the proportion of productively infected cells decayed from 4-10 days as more cells establish latency by epigenetic mechanisms (Fig. 2*A*) [25]. In contrast, Jurkat cells treated with IACS-9571 throughout the course of infection produced a significantly greater proportion of productively infected cells, such that ∼50% of infected cells expressed GFP 4 days post-infection (Fig. 2*A*, IACS-9571). Throughout the course of treatment, negligible effects upon cell viability were observed (Fig. 2*B*). These observations indicate that the TRIM24 bromodomain inhibitor dissuades establishment of immediate latency in newly infected cells and forces productive expression of viral RNAs from the 5’ LTR.

**Figure 2.**
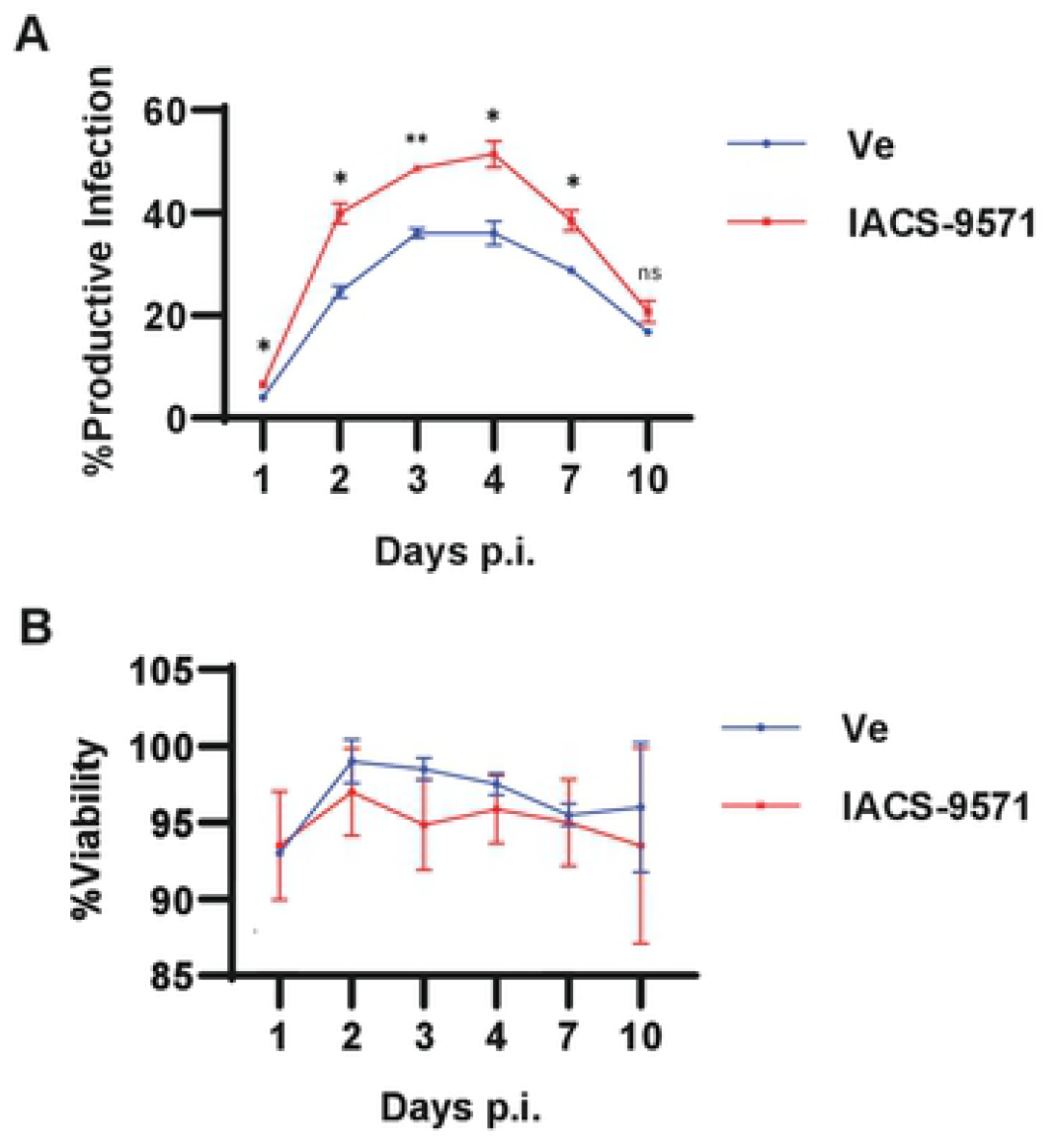
IACS-9571 enforces productive HIV-1 upon infection. **Panel A:** Wildtype Jurkat T cells infected with RGH were treated with or without 10 μM IACS-9571 throughout the course of infection. Flow cytometry was performed on the indicated day post-infection (p.i.). Shown are the results of duplicate experiments, and error bars represent standard deviation. **Panel B:** Cell viability was determined at each time post infection. Results of two measurements are shown, with error bars representing standard deviation.

### IACS-9571 produces synergistic effects with T cell signaling agonists

HIV-1 transcription is activated by multiple transcriptional activators under the control of parallel immune cell signaling pathways stimulated by second messengers and interactions produced by activation of membrane associated protein kinases regulated by engagement of the T cell receptor and various cytokines, including TNF-α and IL2 [12] [8]. These signaling pathways cause activation of multiple transcriptional activators bound to the LTR enhancer region, including NF-κB, AP1, GABP/ Ets, NFAT, and USF1/2-TFII-I (RBF-2) [26][5]. To examine how IACS-9571 might influence responses to these various signaling mechanisms, we measured the effect of this compound on HIV-1 expression in combination with agonists of T cell signaling pathways. Cell line KB1 bearing the integrated LTR-luciferase mini-virus reporter (Fig. 1*A*) were treated with 10 μM IACS-9571 and luciferase expression was measured over the next 24 hours. At this concentration, IACS-9571 on its own produces a modest, but significant ∼2-fold induction of luciferase expression at 6 hours (Fig. 3*A*). However, we found this concentration of IACS-9571 produced a synergistic effect for activation of HIV-1 expression in combination with every T cell signaling agonist we examined. This includes PMA (Fig. 3*B*), which causes activation of multiple factors (NF-κB, AP1, GABP/ Ets); Ionomycin (Fig. 3*C*), which activates NFAT; TNF-α (Fig. 3*E*), which stimulates NF-κB and AP1; and PEP005 (Fig. 3*F*), a latency reversing agent which acts through NF-κB. In contrast, co-treatment of the reporter cells with a combination of IACS-9571 and the HDAC inhibitor SAHA did not produce a synergistic effect on LTR-luciferase expression (Fig. 3*G*). These observations are consistent with previous experiments demonstrating synergy between various latency reversing agents that function through independent mechanisms [7], and suggest that inhibition of the TRIM24 bromodomain likely does not cause reactivation of HIV-1 expression by acting as an upstream agonist for activators bound to the LTR enhancer region.

**Figure 3.**
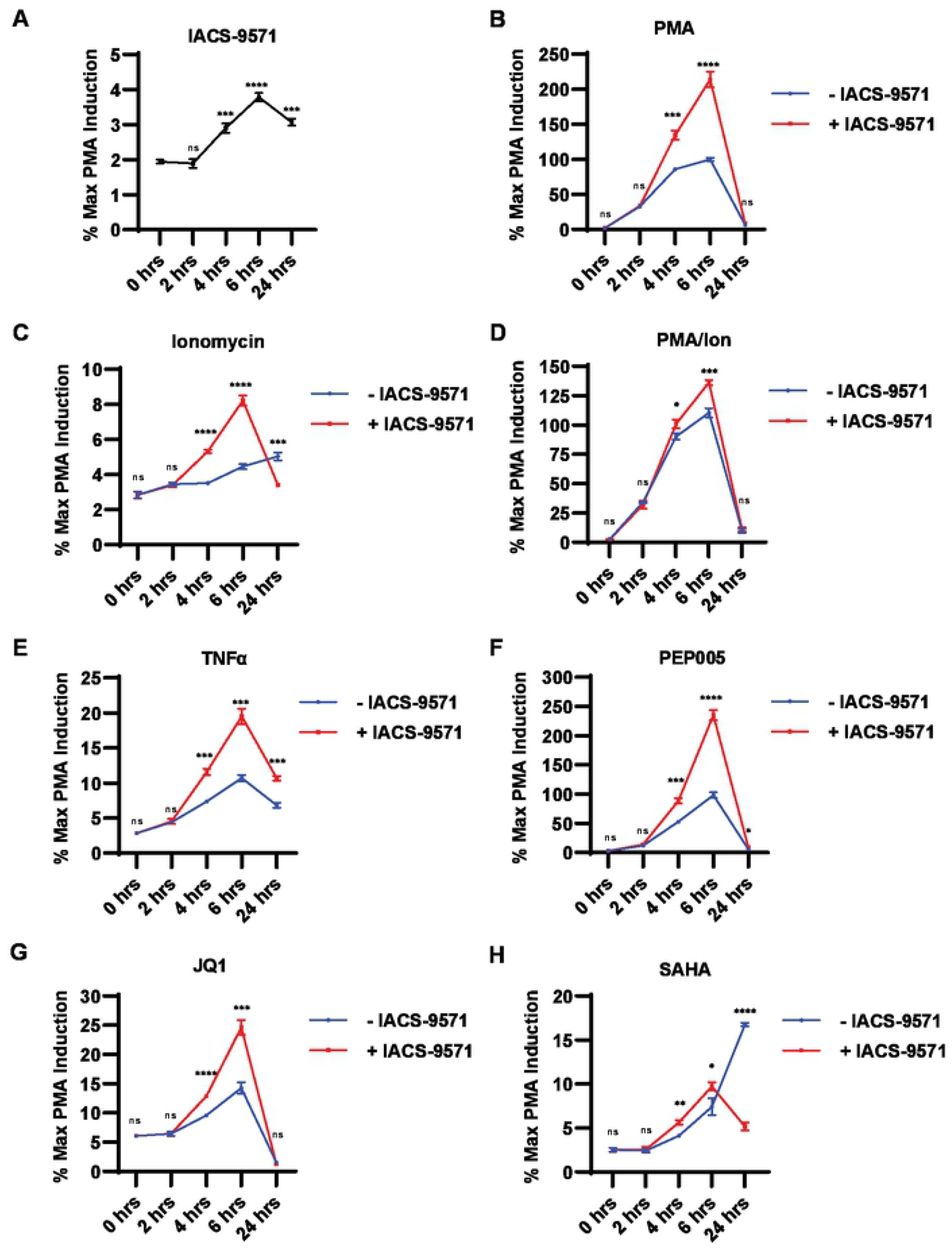
IACS-9571 produces activation in combination with T cell signaling agonists. **Panels A - H:** KB1 cells were treated with the indicated latency reversing agent, with or without 10 μM IACS-9571. Luciferase assays were performed at the indicated times. Results of three determinations are displayed, with error bars representing standard deviation. Concentration of agonist used were 20 nM PMA, 1 μM ionomycin, 10 ng/μL TNFα, 20 nM PEP005, 10 μM JQ1, 10 μM SAHA.

### TRIM24 is necessary for activation of HIV-1 expression, including by IACS-9571

The TRIM24 bromodomain has similar structure to that of another protein designated BRPF1, and IACS-9571 was initially shown to bind both of these bromodomains with similar affinity [24]. Consequently, it is possible that the effects we observe for IACS-9571 on the HIV-1 LTR may be mediated by a factor other than TRIM24, including possibly BRPF1 [24]. To examine this possibility, we compared the effect of IACS-9571 on reactivation of HIV-1 provirus in the KB1 Jurkat cell line bearing the HIV-1 Luciferase reporter provirus (Fig. 1A), and a derivative line bearing a CRISPR/ Cas9-mediated gene disruption of *TRIM24*, designated RTK3 (Fig. 4A, RTK3). As reported above, we observe dose-dependent induction of HIV-1 expression in WT cells treated with IACS-9571 (Fig. 4*B*, KB1), but this effect is completely inhibited in RTK3 cells bearing the *TRIM24* knockout (Fig. 4*B*, RTK3). A similar effect was observed in cells treated with both IACS-9571 and PMA (Fig. 4*C*, KB1). Here, we again observed a synergistic effect of this treatment on HIV-1 expression in WT KB1 cells, but reactivation is significantly inhibited in the *TRIM24* knockout line (RTK3), with only slight activation at the highest concentrations of IACS-9571. These results indicate that the effect of IACS-9571 on HIV-1 expression is largely dependent on the function of TRIM24.

**Figure 4.**
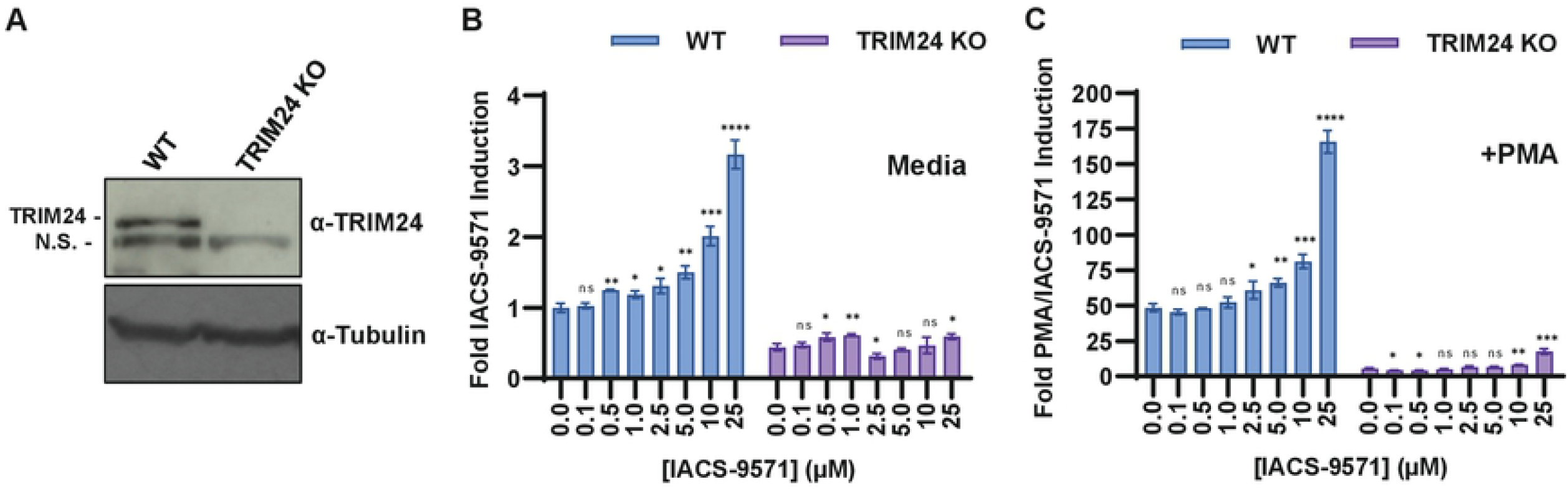
Loss of TRIM24 prevents activation HIV-1 expression in response to IACS-9571. **Panel A:** CRISPR-Cas9 was used to produce a *TRIM24* knockout (KO, lane 2) in the KB1 Jurkat cell line (lane 1). Lysates were analyzed by immunoblotting using antibodies against TRIM24 (top) or tubulin (bottom). **Panel B:** Wildtype and *TRIM24* KO cells were treated with the indicated concentration of IACS-9571 for 4 hrs prior to luciferase assay. Assays were performed in triplicate, with error bars representing standard deviation. **Panel C:** Same as B, but cells were co-treated with 20 nM PMA. Shown are the results of triplicate measurements, error bars represent standard deviation.

### PROTAC-induced degradation of TRIM24 inhibits HIV-1 expression

To examine the requirement of TRIM24 for induction of HIV-1 we used a Von Hippel-Lindau (VHL)-engaging functional degrader of TRIM24, designated dTRIM24 [27]. This chemical derivative is comprised of IACS-9571 conjugated to the VHL binding ligand VL-269 to produce proteolysis-targeting chimeric compound (PROTAC), which promote degradation of target proteins by forcing interaction with the VHL ubiquitin ligase [28]. Treatment of cells with dTRIM24 conjugate causes degradation of TRIM24 in Jurkat cells at concentrations between 0.5-5 mM after 24 hours, as determined by immunoblotting (Fig. 5B, lanes 2-5). As with previous observations, dTRIM24 mediated degradation is lost at higher concentrations (Fig. 5B, lane 6), an effect that is proposed to be the result of the molecule favoring binary interaction over ternary complex formation [27]. Having verified dTRIM24 concentrations that produce its degradation, we measured induction of HIV-1 provirus in these treated cells by analyzing dsRed expression in a cell line bearing a mini-dual HIV reporter [25], 20 hours following treatment with PMA. Consistent with our previous results indicating that TRIM24 is required for reactivation of latent HIV-1 [18] we found that cells treated with concentrations of dTRIM24 that reduce TRIM24 protein levels also displayed a corresponding decrease in expression of the 5’ LTR dsRed reporter in these cells (Fig. 5*C*). In contrast, treatment of this same cell line with IACS-9571 caused significant induction of HIV-1 dsRed expression (Fig. S2). These results demonstrate that the TRIM24 bromodomain inhibitor IACS-9571 produces an opposite effect on HIV-1 transcription than does depletion of this co-factor.

**Figure 5.**
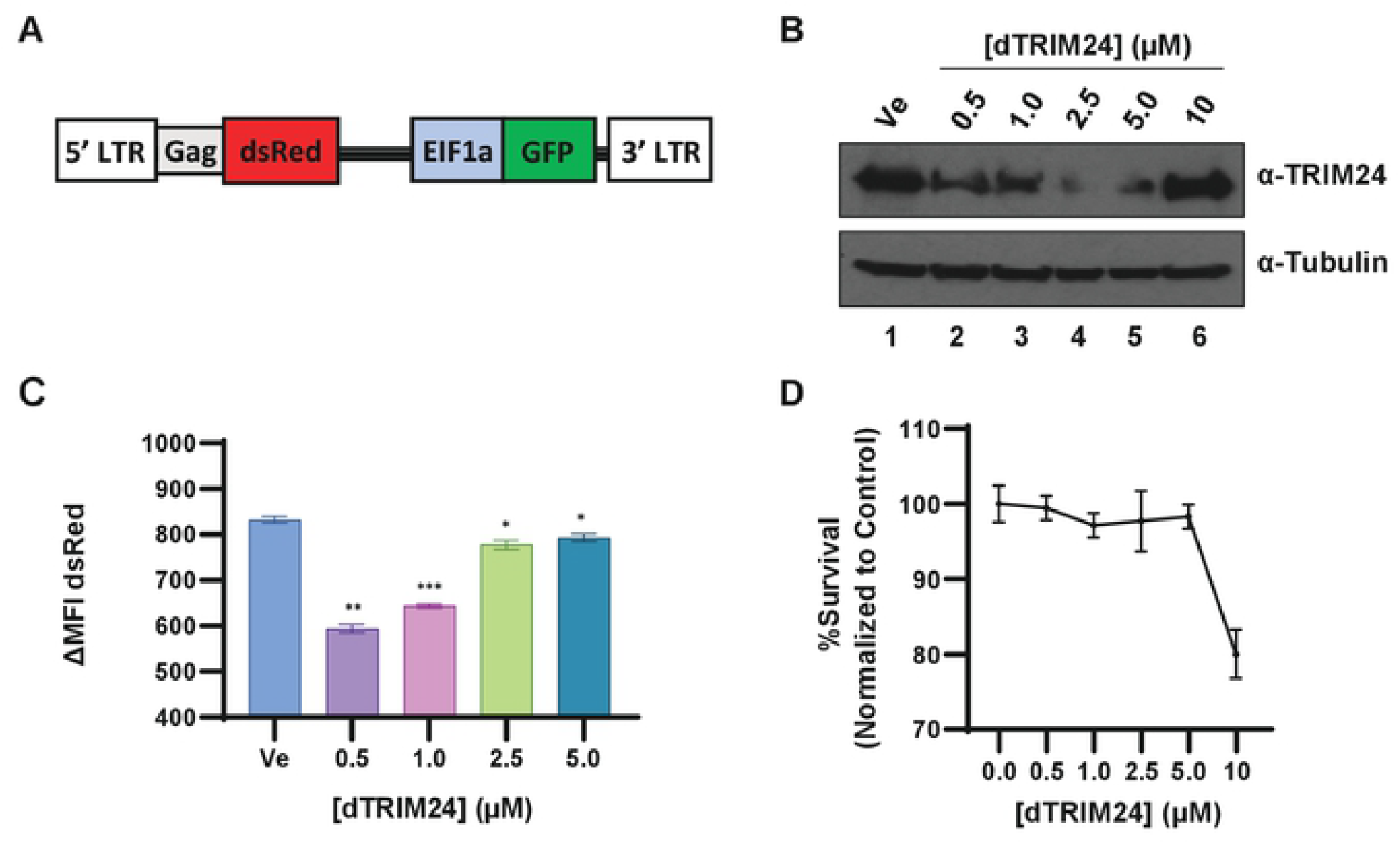
Induced degradation of TRIM24 inhibits HIV-1 expression. **Panel A:** Schematic representation of HIV-1 mini-virus integrated in the Clone #11 Jurkat Tat cell line, where dsRed is expressed from the 5’ LTR, while GFP is driven by the constitutive EIF1α promoter. **Panel B:** Clone #11 cells were treated with the indicated concentration of dTRIM24 or left untreated (Ve) for 24 hrs. Immunoblots were performed on whole cell lysates, with antibodies against TRIM24 (top) or Tubulin (bottom). **Panel C:** Following 4 hr pre-treatment with the indicated concentration of dTRIM24, Clone #11 cells were stimulated with 40 nM PMA for 20 hrs and analyzed by flow cytometry. Assays were performed in duplicate, with error bars representing standard deviation displayed. **Panel D:** Cell viability was measured in cells as treated in panel C. Results are an average of two determinations, and error bars represent standard deviation.

### The C-terminal TRIM24 bromodomain is dispensable for activation of HIV-1 expression

Because the IACS-9571 TRIM24 bromodomain inhibitor produces the opposite effect as dTRIM24, we examined whether the C-terminal bromodomain of TRIM24 was necessary for activation of HIV-1 expression. To this end, we co-transfected HEK293T cells with vectors expressing WT TRIM24, or a series of TRIM24 mutants bearing ORF deletions and point mutations in the C-terminal PHD-bromodomain motifs (Fig. 6*A, B)* in combination with an HIV-1 LTR-luciferase reporter gene. Using this assay, we have previously demonstrated that the effect of TRIM24 for activation of HIV-1 expression does not require the viral transactivator TAT [18]. As with previous experiments, co-transfection of WT TRIM24 expression plasmid causes ∼3 fold stimulation of the HIV-1 luciferase expression (Fig. 6*B*, TRIM24). Interestingly, this effect of TRIM24 does not require the C-terminal PHD-bromodomain region, as a deletion lacking the entire C-terminus (Fig. 6*B*, ΔC-Terminus) causes a similar effect as WT TRIM24. Similarly, derivatives bearing amino acid substitutions in the C-terminal PHD-bromodomain motifs (F979A, N980A, C840W) that disrupt chromatin binding cause similar levels of HIV-l luciferase expression as WT (Fig. 6*B*). In contrast, deletion of the complete N-terminus or the coiled-coil motif (Fig. 6C) prevented stimulation of HIV-1 luciferase activity by TRIM24 (Fig. 6*D*). A mutant with deletion of the BB2 motif was less effective for stimulation of HIV-1 expression (Fig. 6D), but we note that this protein is expressed at significantly lower levels than the other TRIM24 derivatives (Panel C). These results indicate that the bromodomain motif is not required for the effect of TRIM24 for reactivation of HIV-1 transcription.

**Figure 6.**
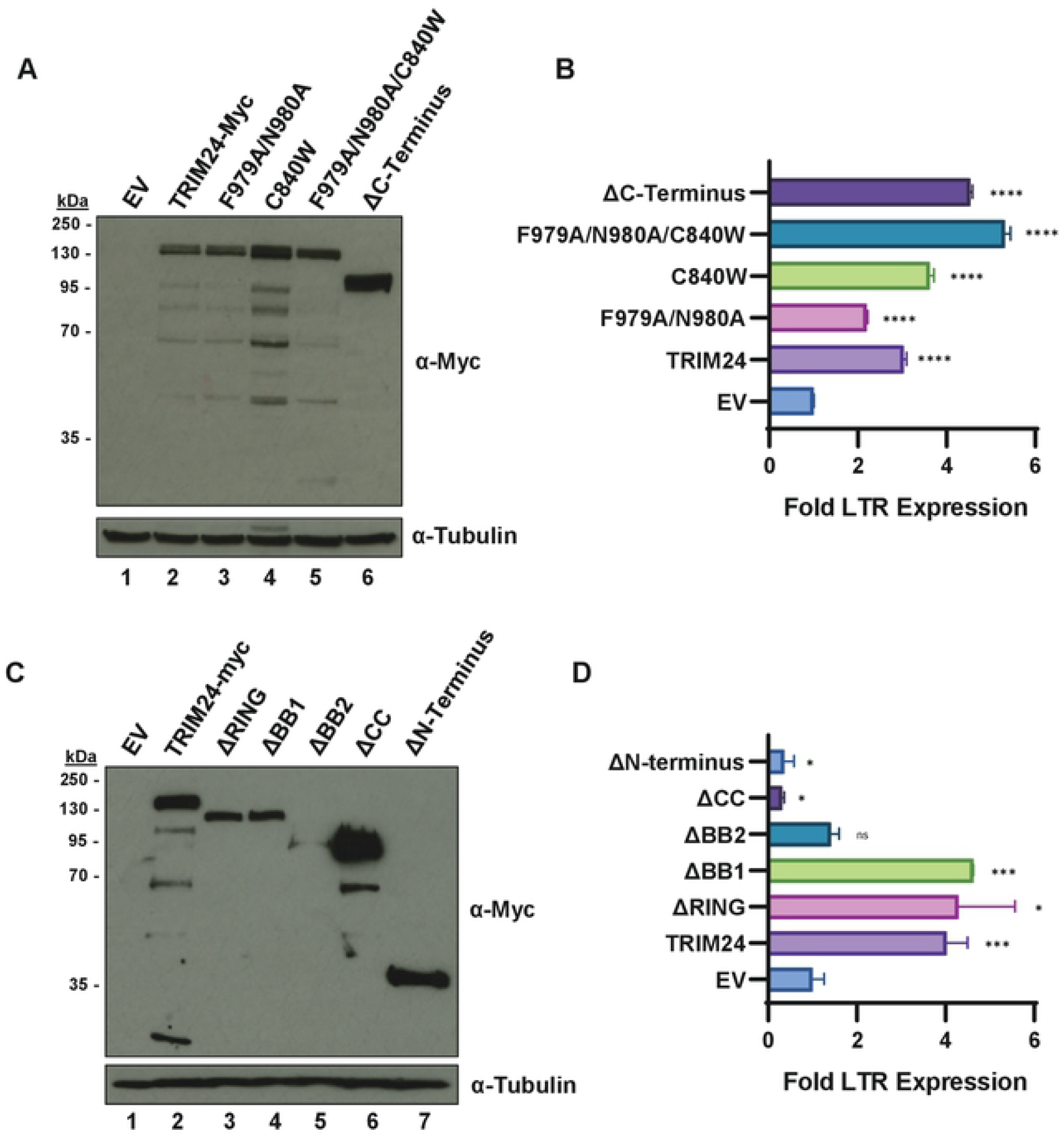
TRIM24 chromatin binding is dispensable for activation of HIV-1 transcription. **Panel A, C:** Two days post-transfection of HEK293T cells with the indicated construct, whole cell lysates were collected and immunoblotted using antibodies against Myc (top) or tubulin (bottom). **Panel B, D:** HEK293T cells were co-transfected with LTR-Luciferase reporter construct and the indicated expression vector. Two days post-transfection, LTR expression was analyzed by luciferase assay. Results are the average of three determinations with standard deviation indicated.

### TRIM24 LTR occupancy is enhanced in response to IACS-9571 treatment

We have previously shown that TRIM24 is associated with the HIV-1 LTR at the RBE3 and RBE1 elements as mediated by direct interaction with the transcription factor TFII-I [18]. The TRIM24 C-terminal tandem bromodomain – plant homeodomain (PHD) is capable of binding histones with preference for H3K23ac and H3K4me0 marks, respectively [21]. Consequently, because TRIM24 is presumed to be predominantly recruited to chromatin *via* interactions mediated by the C-terminal bromo/ PHD domains, we examined the effect of bromodomain inhibition on recruitment of TRIM24 to the HIV-1 LTR. To examine this, we expressed Flag tagged TRIM24 in Jurkat cells harboring an HIV-1 reporter virus (Fig. 7*A*, lanes 2 and 3). Using ChIP-qPCR, we observed significantly enhanced interaction of TRIM24 with the LTR in cells treated with IACS-9571 (Fig. 7*B*). This observation is consistent with our previous results indicating that TRIM24 is a limiting co-factor for activation of HIV-1 expression [18], and suggests that inhibition of the TRIM24 bromodomain may inhibit global interaction of this factor with chromatin, thereby increasing the pool of TRIM24 available for recruitment by TFII-I bound to the HIV-1 LTR.

**Figure 7.**
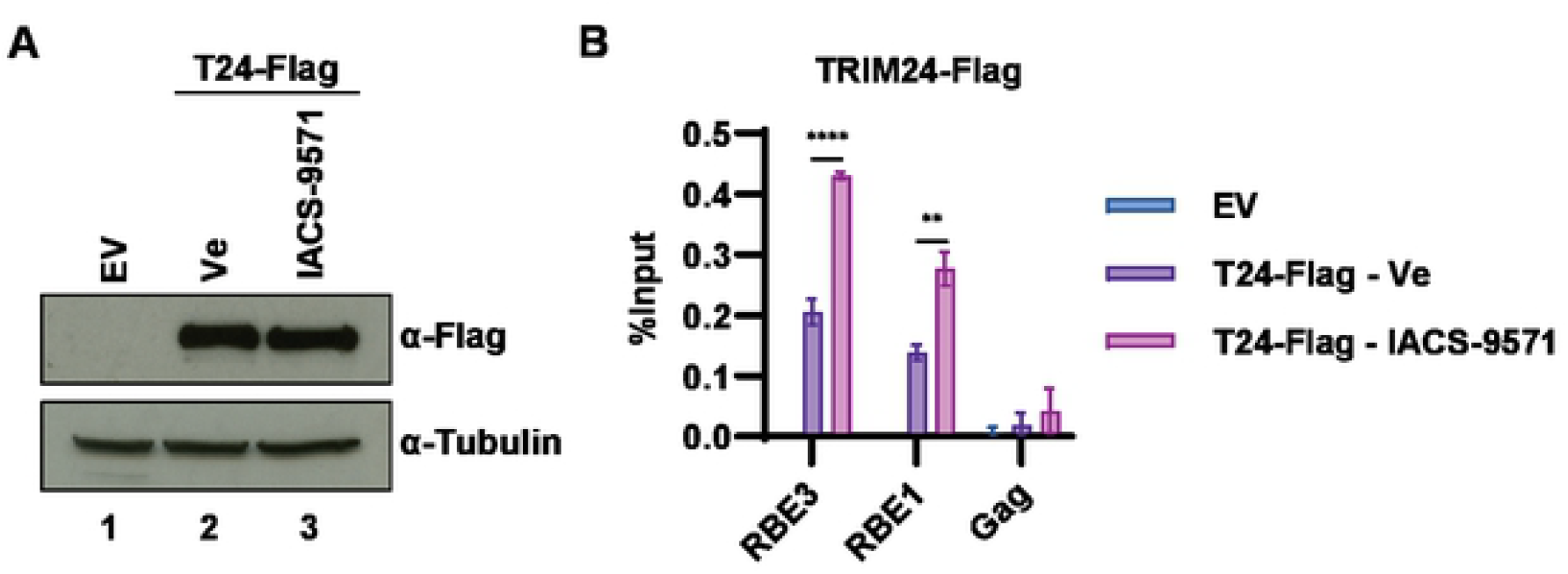
Association of TRIM24 with the HIV-1 LTR is enhanced upon IACS-9571 treatment. **Panel A:** KB1 Jurkat cells were transduced with lentivirus expressing Flag tagged TRIM24 (T24-Flag, Lanes 2, 3) or an empty vector (EV, Lane 1). TRIM24-Flag transduced cells were treated with 10 μM IACS-9571 (Lane 3) or left untreated (Ve, Lane 2). Whole cell lysates were immunoblotted using the indicated antibody. **Panel B:** TRIM24-Flag transduced cells were untreated or incubated with 10 μM IACS-9571. Following 4 hrs, ChIP was performed using the indicated antibody. Results are the average of three qPCR determinations with standard deviation shown.

### IACS-9571 stimulates HIV-1 transcriptional elongation

Our previous results indicate that recruitment of TRIM24 to the HIV-1 LTR by TFII-I causes enhanced elongation of transcription by RNA Polymerase II from the viral core promoter [18]. Consequently, we examined whether treatment of cells with IACS-9571, which causes enhanced association of TRIM24 with the HIV-1 LTR also stimulates transcriptional elongation. We found that treatment of the Jurkat KB1 reporter cell line (Fig. 1*A*) with IACS-9571 did not cause enhanced recruitment of RNAPII to the LTR, rather we observe a slight decrease in PolII occupancy at the LTR in otherwise untreated cells (Fig. 8*A*, compare Ve, IACS-9571). Initiation of eukaryotic transcription is associated with phosphorylation of RNAPII at the C-terminal domain (CTD) serine 5, a modification mediated by Cdk7 of TFIIH [29]. As with total RNA Pol II occupancy, we observed slightly less pS5-modified RNAPII following IACS-9571 treatment as compared to untreated cells (Fig. 8*B*). Elongation of transcription from the LTR is stimulated by recruitment of pTEF-b, containing Cdk9, which phosphorylates RNAPII CTD S2, among other factors, to promote transition from a paused to elongating transcription complex [30]. Interestingly, in contrast to RNAPII and CTD pS5, we found that treatment of Jurkat KB1 cells with IACS-9571 cause elevated association of RNAPII pS2 with the LTR as compared to untreated cells (Fig. 8*C*). Additionally, IACS-9571 treatment caused significant enrichment of CDK9 (PTEF-b) at the HIV-1 core promoter (Fig. 8*D*, RBE1). These results are consistent with our previous observations indicating that recruitment of TRIM24 to the HIV-1 LTR causes enhanced recruitment of pTEF-b/ Cdk9, which promotes RNAPII CTD S2 phosphorylation and elongation of transcription [18].

**Figure 8.**
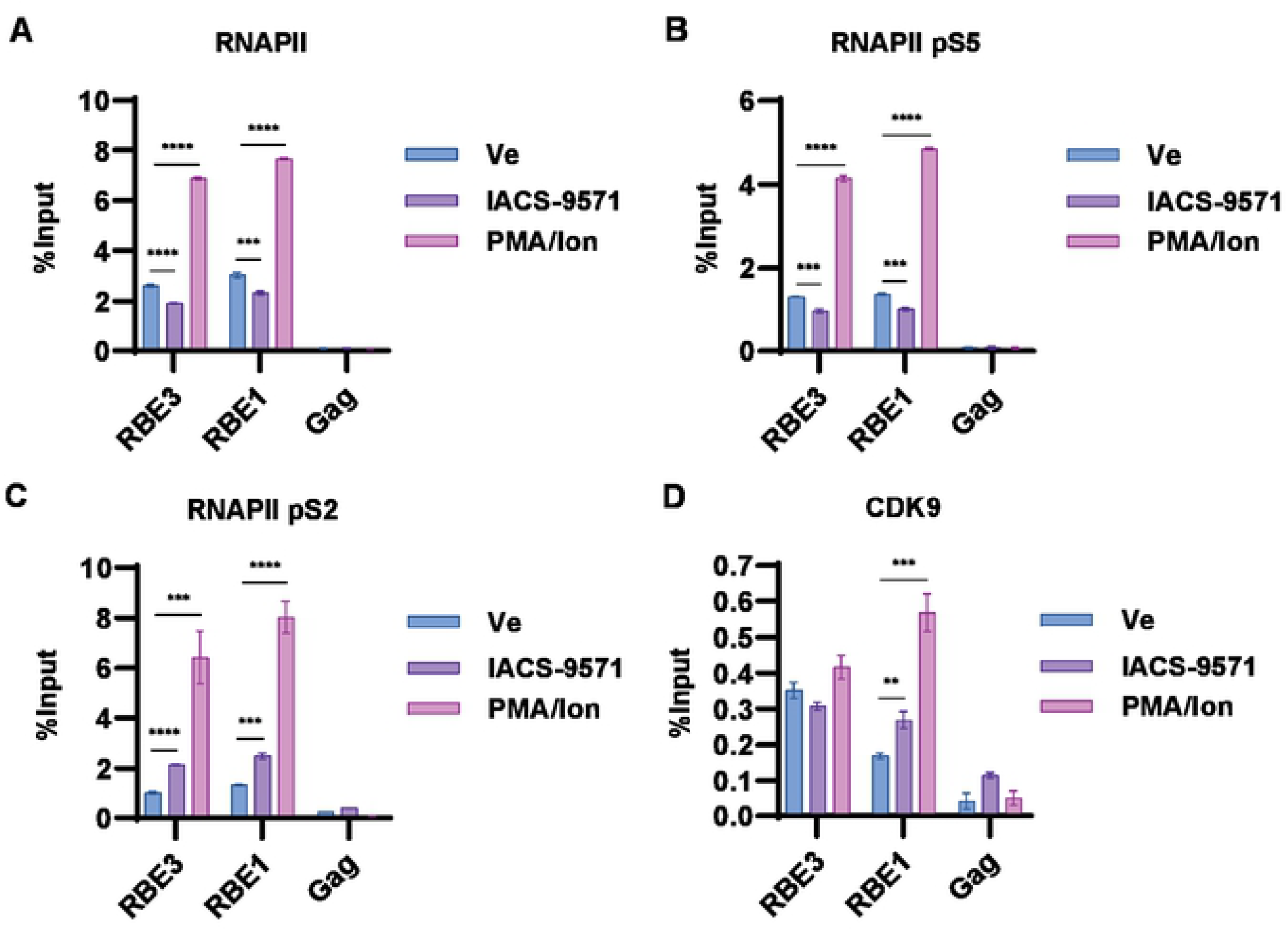
IACS-9571 stimulates elongation from the HIV-1 LTR. **Panel A-D:** KB1 Jurkat cells were treated with 10 μM IACS-9571, 20 nM PMA/ 1 μM Ionomycin, or the equivalent amount of DMSO (Ve). After 4 hrs treatment, ChIP was performed using the indicated antibody. Displayed are the average of three qPCR determinations with error bars representing standard deviation.

Results shown above (Fig. 3) indicate that IACS-9571 causes synergistic effects of HIV-1 provirus expression in combination with T cell signaling agonists. We note that treatment with PMA/ Ionomycin or PEP005, agonists of pathways stimulated by the T cell receptor [31], cause significantly enhanced recruitment of RNAPII (Fig. 8A) and phosphorylation of CTD S5 (Fig. 8B) to the HIV-1 LTR, indicating that activation of factors regulated by T cell signaling have a significant effect on recruitment of RNA Pol II to the LTR promoter for promotion of transcriptional initiation. Co-treatment of cells with IACS-9571 with PMA/ Ionomycin or PEP005 causes corresponding elevated association of RNAPII, CTD pS5 and pS2 with the LTR (Fig8 A, B, C) which is consistent with the synergistic effect on HIV-1 reporter expression produced by the combination of these treatments.

### Effect of IACS-9571 on HIV-1 provirus expression in CD4+T-cells from individuals on ART

To determine whether IACS-9571 was capable of affecting expression of HIV-1 provirus in primary CD4^+^ lymphocytes, we examined the effect of treatment on CD4^+^ T-cells isolated from individuals with HIV-1 on antiretroviral therapy. We found that treatment with 10 μM IACS-9571 on its own did not cause expression of HIV-1 mRNA in any of the cell samples from 6 participants examined (Fig. 9, compare Ve and IACS-9571). Consistent with previous observations we found that 15 nM PEP005, a PKC agonist [7], caused a significant increase in viral transcription in cells from two of the participants (Fig. 9*A*, 9*E*, BC003 and BC008), and a more modest effect on a third participant (Fig. 9D, BC006). Interestingly, with cells from all of the participant samples, co-treatment with IACS-9571 and PEP005 caused significant induction of HIV-1 mRNA (Fig. 9), at concentrations that do not affect cell viability (Fig. S3), a result that is consistent with the synergistic effects these agents have on induction of HIV-1 reporter expression in cell lines (Fig. 3).

**Figure 9.**
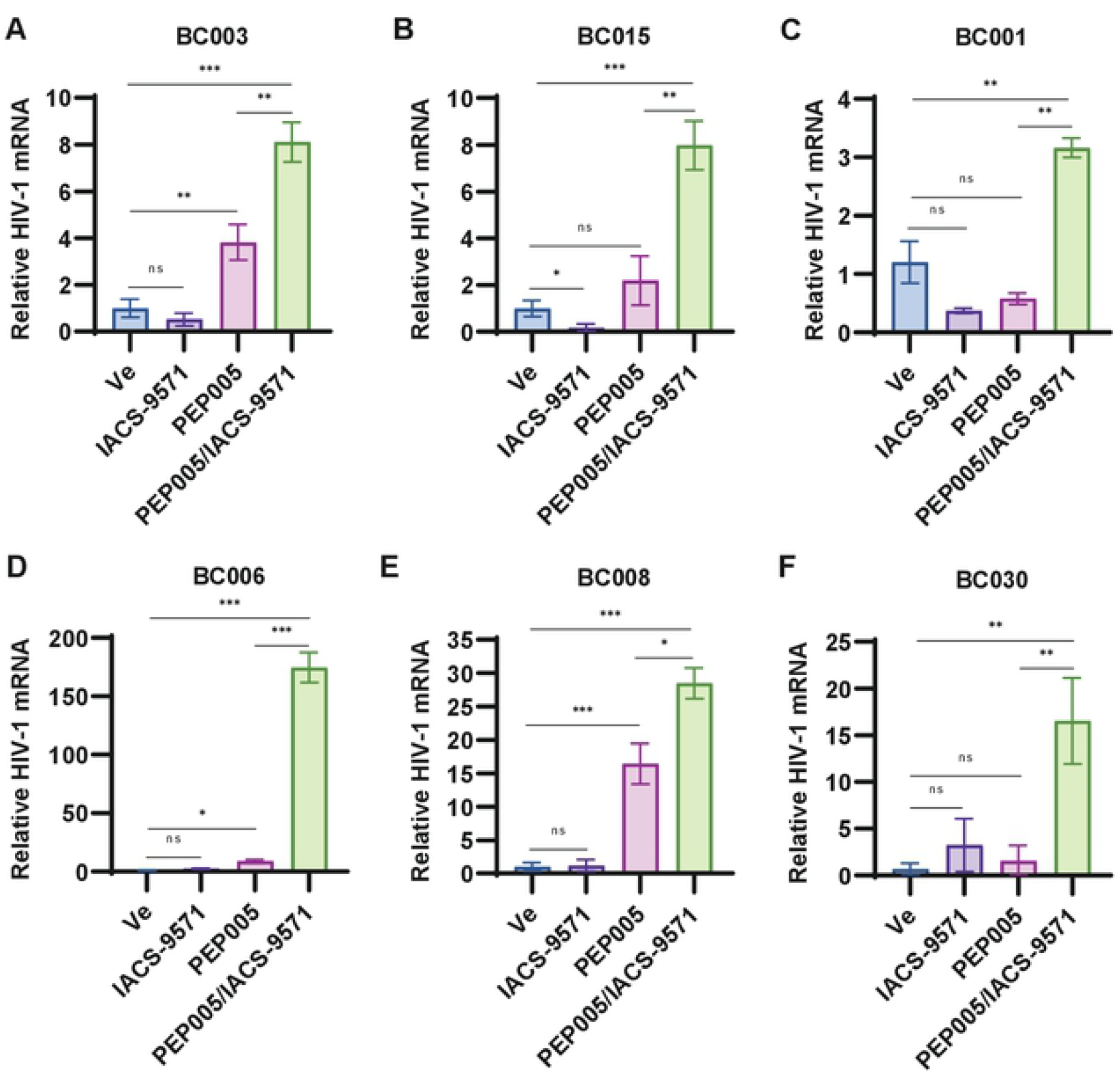
IACS-9571 and PEP005 synergistically activate HIV-1 expression in PBMCs. **aPnel A - F:** CD4^+^ PBMCs isolated from HIV-1 patients on ART were treated with DMSO (Ve), 10 μM IACS-9571, 15 nM PEP005, or with 10 μM IACS-9571 and 15 nM PEP005 for 20 hrs. Following incubation, intracellular RNA was extracted and subject to RT-PCR using oligos targeting HIV-1 mRNA. The results of three RT-PCR determinations are displayed with standard deviation depicted as error bars.

### IACS-9571 causes partial inhibition of T cell activation

An important consideration for development of latency reversing agents as potential therapies is that treatment should produce robust and broad induction of HIV-1 expression without simultaneously causing global T cell activation. To determine the potential effect of IACS-9571 on T cell activation, we examined expression of IL2 and CD69 mRNAs in cells from participants using Q-RT-PCR. Consistent with previous results [32], treatment of CD4^+^ cells from participants with HIV-1 on ART with the PKC agonist PEP005 caused a significant increase in IL2 mRNA (Fig. 10). In contrast, treatment with IACS-9571 on its own did not cause elevation of IL2 mRNA levels. Moreover, we observed that treatment with IACS-9571 in combination with PEP005 resulted in suppression of IL2 mRNA induction (Figure 10). Similar results were obtained for expression of CD69, a cell surface marker associated with T cell activation (Fig. S4). Furthermore, we observe a comparable effect on IL2 and CD69 expression in Jurkat T cells treated with the combination of PMA and IACS-9571 (Fig. S5). These results indicate that IACS-9571 must affect expression of the IL2 and CD69 genes by a different mechanism than the HIV-1 LTR. Importantly as well, no previous compound with latency reversing activity was found to produce opposite effects for T cell activation and induction of HIV-1 expression [7].

**Figure 10.**
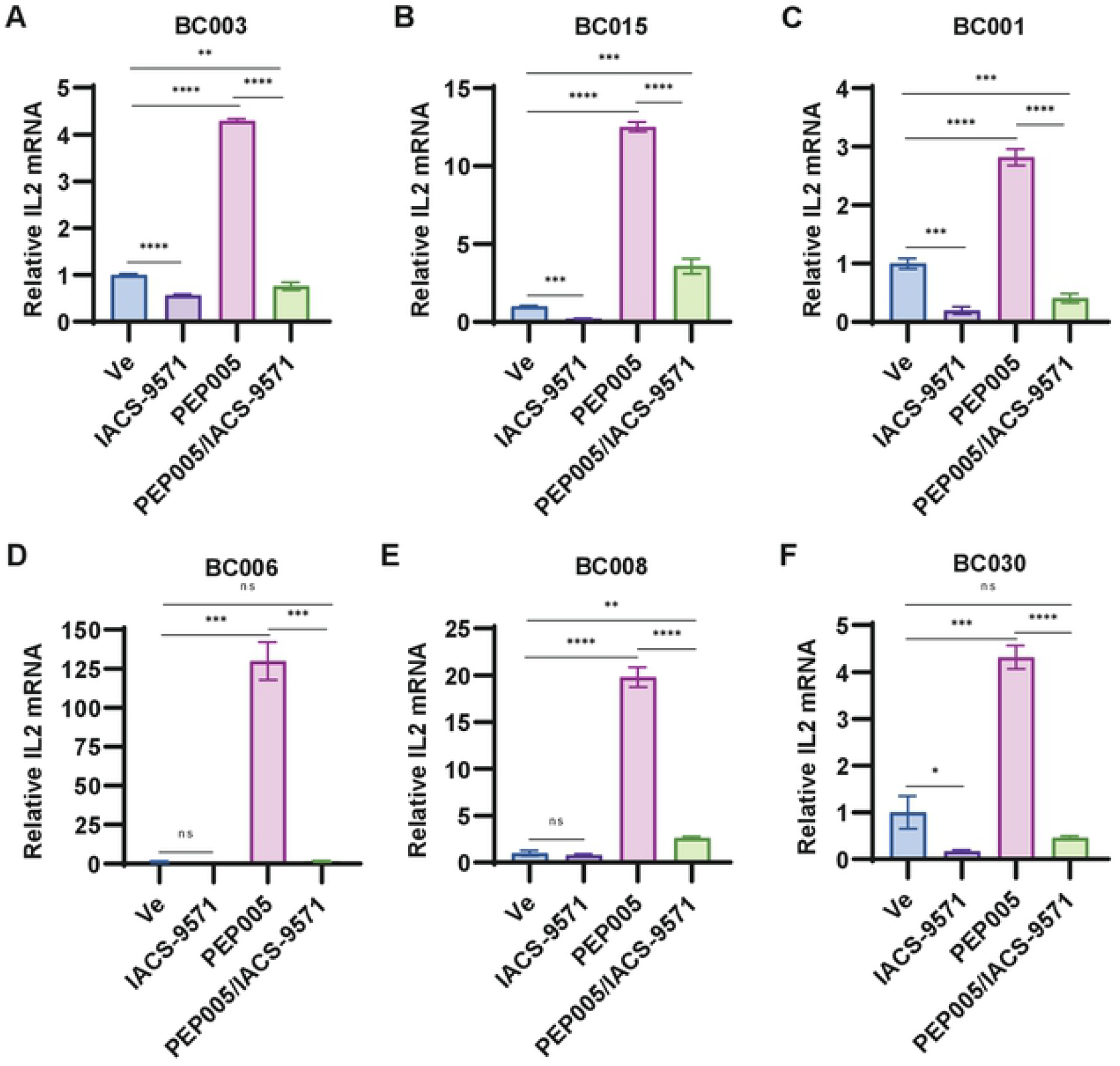
IACS-9571 does not cause T cell activation. **Panel A - F:** CD4^+^ PBMCs isolated from ART administered HIV-1 patients were treated with DMSO (Ve), 10 μM IACS-9571, 15 nM PEP005, or with 10 μM IACS-9571 and 15 nM PEP005 for 20 hrs. Following treatment, intracellular RNA was extracted and subject to RT-PCR using oligos that target IL2 mRNA. RT-PCR was performed in triplicate with error bars representing standard deviation.

## Discussion

TRIM24 was initially identified as a co-activator of transcription for various nuclear receptors [33]. This factor has E3 ubiquitin ligase activity, conferred by the N-terminal RING motif, and interacts with chromatin *via* the C-terminal PHD-bromodomain through binding of histone H3K40K23ac [20]. Despite its discovery as a transcriptional co-activator ∼25 years ago [33], a mechanism for its function in transcriptional regulation had not been identified. We recently discovered that TRIM24 is recruited to the HIV-1 LTR by interaction with TFII-I, which promotes elongation of transcription in stimulated T cells through enhanced recruitment of pTEF-b/ Cdk9 and phosphorylation of RNAPII CTD S2 [18]. Consequently, we were surprised that the TRIM24 bromodomain inhibitor IACS-9571 caused elevated expression of HIV-1 provirus in otherwise untreated Jurkat T cells. This effect was accompanied by enhanced association of TRIM24 with HIV-1 LTR promoter. Because TRIM24 seems to be a limiting factor for reactivation of HIV-1 transcription [18], our observations suggest that IACS-9571 may inhibit global interaction of this factor with chromatin and consequently increasing opportunity for recruitment to the LTR by interaction with TFII-I. Consistent with previous observations we found that forced degradation of TRIM24 with the dTRIM24 proteolysis-targeting chimeric compound (PROTAC) inhibited HIV-1 provirus expression at concentrations where it caused reduction of TRIM24 protein levels. These observations implicate TRIM24 as an important potential target for therapies to modulate expression of latent HIV-1 provirus in patients on antiretroviral therapy. IACS-9571 causes reactivation of HIV-1 provirus at similar concentrations, and as effectively, as most previously characterized latency reversing agents [7]. Conversely, the bifunctional compound dTRIM24, designed to promote degradation of TRIM24 through a VHL E3-mediated mechanism [24] [27], inhibits reactivation of HIV-1 provirus expression, which indicates that inhibitors of full TRIM24 function, or at least interaction with TFII-I on the LTR would represent potential latency promoting agents.

TRIM24 confers both positive and negative effects on gene expression. In RNA-seq analysis of *TRIM24* knockout cells we observed differential effects on nearly 2000 genes compared to WT, with approximate equal representation of up and down regulated effects on specific gene expression [18]. Details regarding mechanism for the repressive effect of TRIM24 for specific genes is limited, but was reported to involve inhibition of transcriptional activation by the retinoic acid receptor [34] and Smad4 [35] by effects that appear to involve ubiquitylation of these factors. The observation that IACS-9571 reactivates HIV-1 provirus suggests that, in unstimulated cells, TRIM24 may inhibit one or more factors bound to the 5’ LTR to suppress basal expression from the viral promoter. In this view, it is possible that TRIM24 function for repression or activation of HIV-1 expression is altered by T cell signaling mechanisms. TRIM24 is recruited to the LTR by interaction with TFII-I [18], a protein that functions for both repression and activation of HIV-1 provirus expression [31][16][26]. Consequently, it is possible that the divergent effects of factors bound to the conserved RBE1 and RBE3 elements on the LTR may be mediated by differentially regulated functions of TRIM24.

Alternatively, TRIM24 may not have a direct inhibitory effect on transcriptional activation of the HIV-1 LTR, but rather the latency reversing effect of IACS-9571 may be produced by inhibition of its interaction with H3K40K23ac modified chromatin on cellular genes. This effect might enable recruitment of elevated levels of TRIM24 to the LTR in the cellular population for stimulation of transcriptional elongation. Consistent with this view we observe enhanced levels of TRIM24 associated with the LTR in cells treated with IACS-9571 (Fig. 7). Furthermore, we find that TRIM24 mutant proteins bearing deletion of the C-terminal region spanning the histone binding motifs, or point mutations within those motifs, do not affect the capability of this factor to stimulate HIV-1 transcription (Fig. 6). These observations incidentally suggest that the histone binding function of TRIM24 is not directly required for its effect on stimulating transcriptional elongation from the HIV-1 LTR. However, overall a more detailed understanding of TRIM24 function will be required to elucidate the mechanism for function of IACS-9571 as a latency promoting agent.

Unlike most other previously described HIV-1 latency reversing agents, IACS-9571 inhibits expression of genes associated with T cell activation (IL2 and CD69) in CD4^+^ cells from individuals with HIV-1 on ART, and in Jurkat T cells stimulated with PMA. An important consideration for development of latency reversing agents is their effectiveness for reactivation of HIV-1 provirus without causing global T cell activation [7]. Unchecked activation of T cells can produce a response known as cytokine storm, typified by excess production of proinflammatory cytokines which can cause acute respiratory distress syndrome. Consequently, effects of cytokine storm caused by hyperactivation of T cells contributes significantly to pathology of Covid-19 [36] and influenza [37] infections. Our observation that IACS-9571 inhibits expression of genes associated with T cell activation, but causes activation of HIV-1 expression illustrates a unique capability of this novel latency reversing agent. We note that the BET family bromodomain inhibitor JQ1 also acts as an LRA and causes down-regulation of T cell activation genes [38]. Our results indicate that TRIM24 protein has opposing mechanistic effects for regulation of HIV-1 expression from genes regulating T cell response; these differential effects must be mediated by differential requirement for function of the C-terminal bromodomain. A more detailed understanding of the mechanistic significance of histone H3 K23 acetylation will be required to elucidate the precise effect of IACS-9571 on HIV-1 transcription. Nevertheless, considering the unique effects this compound has on HIV-1 provirus and T cell activation we suggest that IACS-9571, and its derivatives, may prove useful for shock and kill strategies to purge latently infected cells from individuals with HIV-1 on antiretroviral therapy. We found that, although IACS-9571 on its own does not reactivate proviruses in cells from participants with HIV-1, it causes synergistic reactivation of viral expression in combination with PEP005 (Fig. 9). Consequently, we suggest this combination of treatment may prove useful for implementation of possible shock and kill therapy. Accordingly, we note that this compound was previously proposed as a therapy for glioblastomas where TRIM24 is a contributing factor for malignancy [39], and that PEP005 is already in clinical trials for treatment of skin cancers [40].

## Materials and Methods

### Cell and virus culture

Jurkat and HEK293T cells were maintained in culture conditions as previously described [17]. VSV-G pseudotyped viruses were produced by co-transfection of HEK293T cells with viral molecular clone psPAX and pHEF-VSVg as previously described [17]. Peripheral Blood Mononuclear Cells (PBMC) from participants with HIV-1 on ART were isolated from whole blood by density gradient centrifugation using Lymphoprep™ and SepMate™ tubes (StemCell Technologies), and cryopreserved [41]. Upon thawing, PBMCs were cultured in RPMI 10% FCS. Samples from donor participants were collected with written consent in accordance with UBC Human Ethics certificate H16-02474.

### Immunoblotting

Western blotting was performed as previously described [41]. Antibodies were as follows: Tubulin, Abcam ab7291; Flag, Sigma Aldrich F3165; Myc, Santa Cruz sc-40; TRIM24, Proteintech 14208-1-AP; TFII-I, Abcam ab134133; Goat Anti-Rabbit-HRP, Abcam ab6721; Goat Anti-Mouse-HRP, Pierce 1858413.

### Chromatin Immunoprecipitation

Jurkat Tat KB001 cells were fixed with 1% formaldehyde (Sigma-Aldrich) for 10 min at room temperature (3×10^7^ cells/IP). 125 mM glycine was added for 5 min to quench cross-linking, and cells were subsequently washed with PBS on ice. Cells were lysed in NP-40 Lysis Buffer (0.5% NP-40, 10 mM Tris-HCl pH = 7.8, 3 mM MgCl_2_, 1x PIC, 2.5 mM PMSF) for 15 min on ice. Following sedimentation, nuclei were resuspended in Sonication Buffer (10 mM Tris-HCl pH = 7.8, 10 mM EDTA, 0.5% SDS, 1x PIC, 2.5 mM PMSF) and sonicated using a Covaris S220 Focused-ultrasonicator to produce sheared DNA (2000-200 bp). The soluble chromatin fraction was collected and snap frozen in liquid nitrogen. Chromatin concentrations were normalized among samples and pre-cleared with Protein A/G agarose (Millipore, 100 μL/IP). The chromatin samples were split in two and diluted with IP buffer (10 mM Tris-HCl pH = 8.0, 1.0% triton X-100, 0.1% deoxycholate, 0.1% SDS, 90 mM NaCl, 2 mM EDTA, 1x PIC); samples were immunoprecipitated with the indicated specific antibody or control IgG. Antibodies for ChIP were: TRIM24, Proteintech 14208-1-AP; RNAPII, Abcam ab26721; RNAPII pS5, Abcam ab5408; RNAPII pS2, Abcam #b238146; CDK9, Abcam ab239364; Flag, Sigma Aldrich F3165; Mouse IgG, Santa Cruz sc-2025; Rabbit IgG, Abcam ab1722730. The chromatin/ antibody mixtures were incubated 1 hr at 4^0^C with rotation. Pre-washed Protein A/G agarose beads (40 μL/ IP) were then added and the samples were incubated overnight at 4^0^C with rotation. Bead complexes were washed 3x in Low Salt Wash Buffer (20 mM Tris-HCl pH = 8.0, 0.1% SDS, 1.0% Triton X-100, 2 mM EDTA, 150 mM NaCl, 1x PIC) and 1x with High Salt Wash Buffer (same but with 500 mM NaCl). Elution and crosslink reversal was performed by incubating 4 hrs at 65^0^C in elution buffer (100 mM NaHCO_3_, 1% SDS) supplemented with RNase A. DNA was purified using the QIAQuick PCR purification kit (QIAGEN) and ChIP DNA was analyzed using the Quant Studio 3 Real-Time PCR system (Applied Biosystems). The percent input value of the IgG sample was subtracted from the specific antibody percent input value of the corresponding sample. Oligos used for ChIP-qPCR were: RBE3, Fwd 5’ AGCCGCCTAGCATTTCATC, Rev 5’ CAGCGGAAAGTCCCTTGTAG; RBE1, Fwd 5’ AGTGGCGAGCCCTCAGAT, Rev 5’ AGAGCTCCCAGGCTCAAATC; Gag, Fwd 5’ AGCAGCCATGCAAATGTTA, Rev 5’ AGAGAACCAAGGGGAAGTGA.

### Luciferase reporter assays

Luciferase expression assays from transiently transfected cells were performed as previously described [18]. Transfection of HEK293T cells were performed in 96-well plates seeded with 2×10^4^ cells per well one day prior to transfection. Cells were co-transfected with 10 ng of pGL3 LTR reporter plasmid and 100 ng expression vector; luciferase activity was measured 24 hours post-transfection. For Jurkat luciferase reporter assays, 96-well plates were seeded with 1×10^5^ cells in 100 μL media, and luciferase activity was measured after the indicated time of treatment. Measurements were performed using Superlight™ luciferase reporter Gene Assay Kit (BioAssay Systems) as per the manufacturer’s instructions, and activity was determined by a VictorTM X3 Multilabel Plate Reader.

### Q-*RT-PCR*

Following the indicated treatment, RNA was extracted from cells using an RNeasy Kit (Qiagen) and subsequently analyzed with the Quant Studio 3 Real-Time PCR system (Applied Biosystems) using *Power* SYBR® Green RNA-to-CT™ 1-Step Kit (Thermo Fisher) as per the manufacturer’s instructions. Primers were as follows: IL2, Fwd 5’ AACTCACCAGGATGCTCACA, Rev 5’ GCACTTCCTCCAGAGGTTTGA; CD69, Fwd 5’ TCTTTGCATCCGGAGAGTGGA, Rev 5’ ATTACAGCACACAGGACAGGA; HIV-1 mRNA, Fwd 5’ CTTAGGCATCTCCTATGGCAGGA, Rev 5’ GGATCTGTCTCTGTCTCTCTCTCCACC

### Statistical analyses

Details of statistical analysis are indicated in Figure Legends. Mean is shown with standard deviations. Unpaired sample *t*-tests were performed using GraphPad Prism 9.0.0, and statistical significance is indicated at \**P* < 0.05, \*\**P* < 0.01, \*\*\**P* < 0.001, or \*\*\*\**P* < 0.0001.

## Acknowledgments

We thank Andy Johnson and Justin Wong of the UBC Flow Cytometry Facility for performing FACS analysis as well as for assistance with flow cytometry. We thank the laboratory staff at the BC Centre for Excellence in HIV/AIDS for processing PBMCs from study participants. This research was supported by program project grant F16-01210, from the Canadian Institutes of Health Research (CIHR). PBMC collection from participants with HIV-1 was supported by CIHR project grant PJT-159625 (to ZLB). ZLB is supported by a Michael Smith Health Research BC Scholar Award.

## Author Contributions

Brumme Z.L provided PBMCs from participants with HIV-1 on ART. Horvath R. M. performed all other experiments. Horvath R.M. and Sadowski I. wrote the manuscript.

## Legends to Supplementary Figures

**Figure S1.**
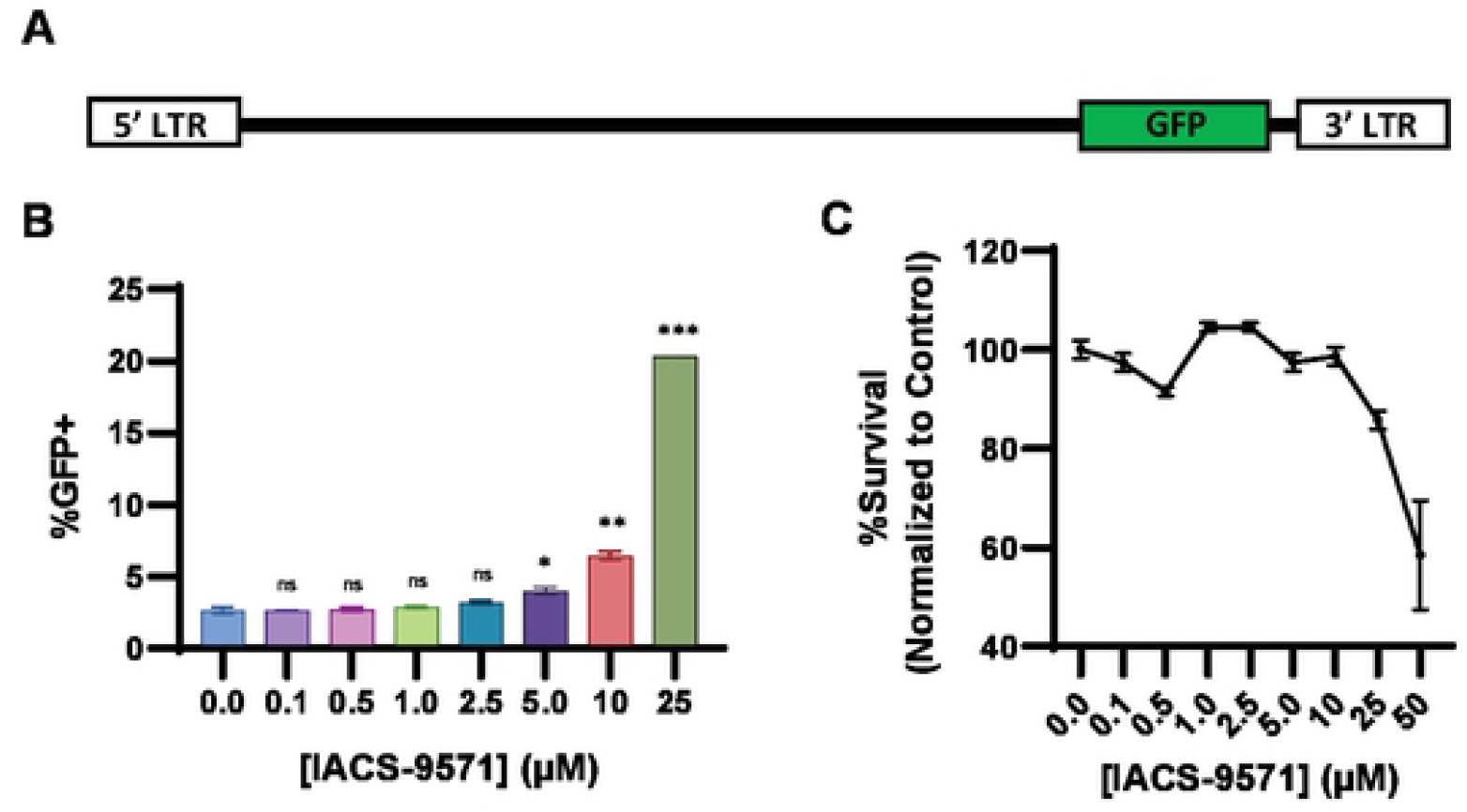
IACS-9571 promotes HIV-1 expression. **Panel A:** Schematic representation of HIV-1 virus integrated in the JLat10.6 cell line. The full-length virus expresses GFP in place of *Nef* with mutations to *Env* rendering it replication incompetent. **Panel B:** JLat10.6 cells were treated with IACS-9571 at indicated concentration for 24 hrs and subsequently analyzed by flow cytometry. Results are the average of two measurements, error bars represent standard deviation. **Panel C:** JLat10.6 cell viability was determined following 24 hrs treatment with the indicated concentration of IACS-9571. Assays were performed in duplicate and error bars depict standard deviation.

**Figure S2.**
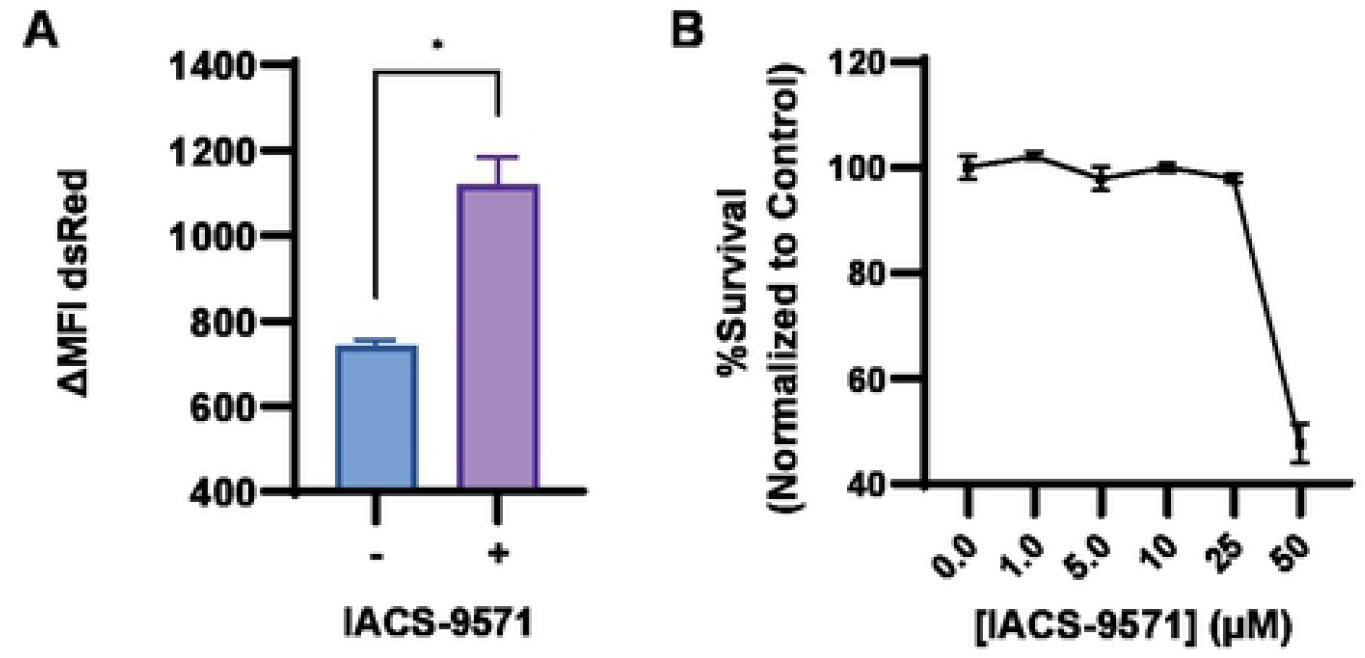
Jurkat Tat Clone #11 is responsive to IACS-9571. **Panel A:** Clone #11 cells were treated with 40 nM PMA or 40 nM PMA and 10 μM IACS-9571. Flow cytometry was performed following 24 hrs incubation. Displayed are the results of duplicate experiments, with error bars representing standard deviation. **Panel B:** Viability of Clone #11 cells treated with 40 nM PMA and the indicated concentration of IACS-9571 was determined following 24 hrs incubation. Displayed are the results of two determinations with error bars depicting standard deviation.

**Figure S3.**
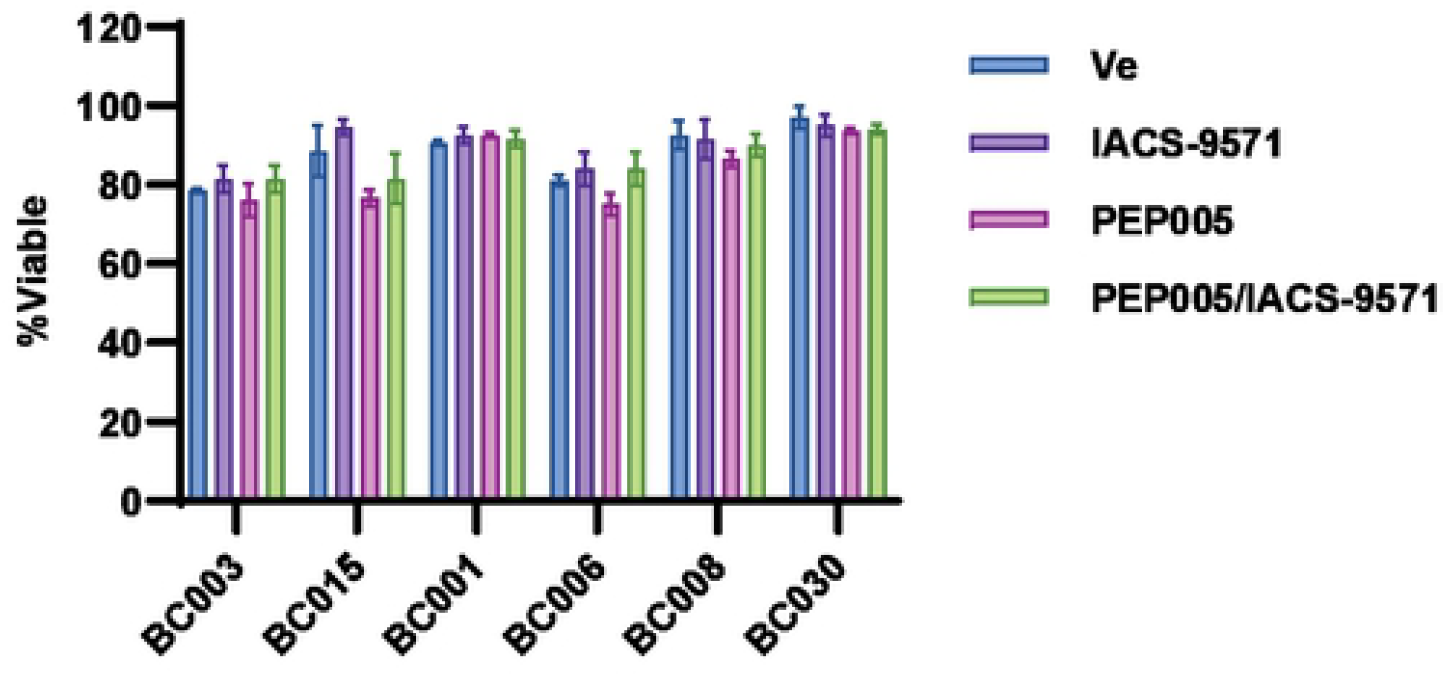
Viability of drug treated HIV-1 patient PBMCs. Cell viability was measured following 20 hrs treatment with DMSO (Ve), 10 μM IACS-9571, 15 nM PEP005, or 10 μM IACS-9571 and 15 nM PEP005. A Bio-Rad TC20 Automated Cell Counter was used to measure viability. Displayed is the average of two independent measurements, and standard deviation is indicated.

**Figure S4.**
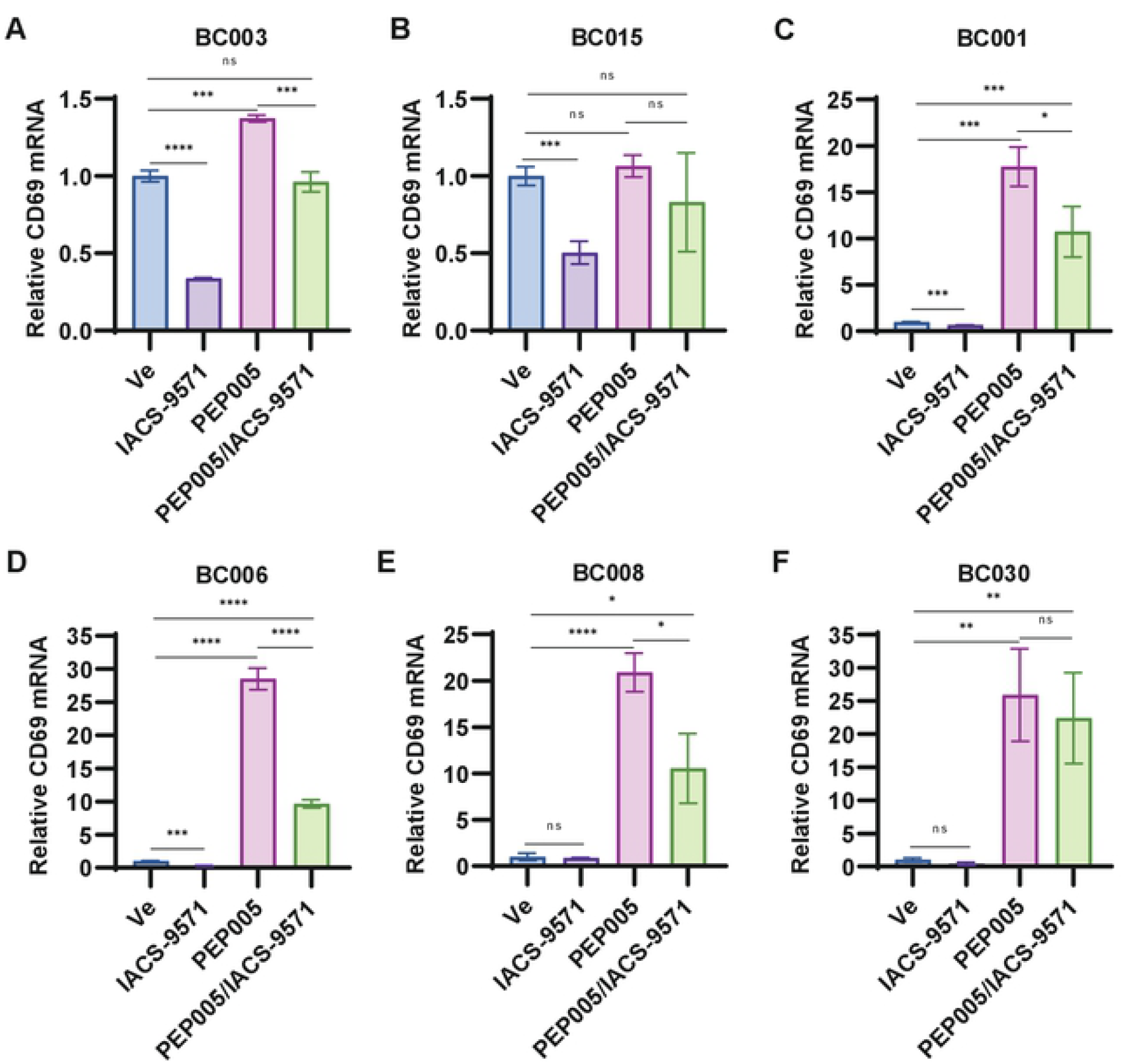
IACS-9571 does not cause T cell activation as measured by CD69 expression. **Panel A - F:** Following 20 hrs treatment with DMSO (Ve), 10 μM IACS-9571, 15 nM PEP005, or 10 μM IACS-9571 and 15 nM PEP005, intracellular RNA was extracted from HIV-1 patient PBMCs. RT-PCR using oligos targeting CD69 mRNA was performed in triplicate with error bars representing standard deviation.

**Figure S5.**
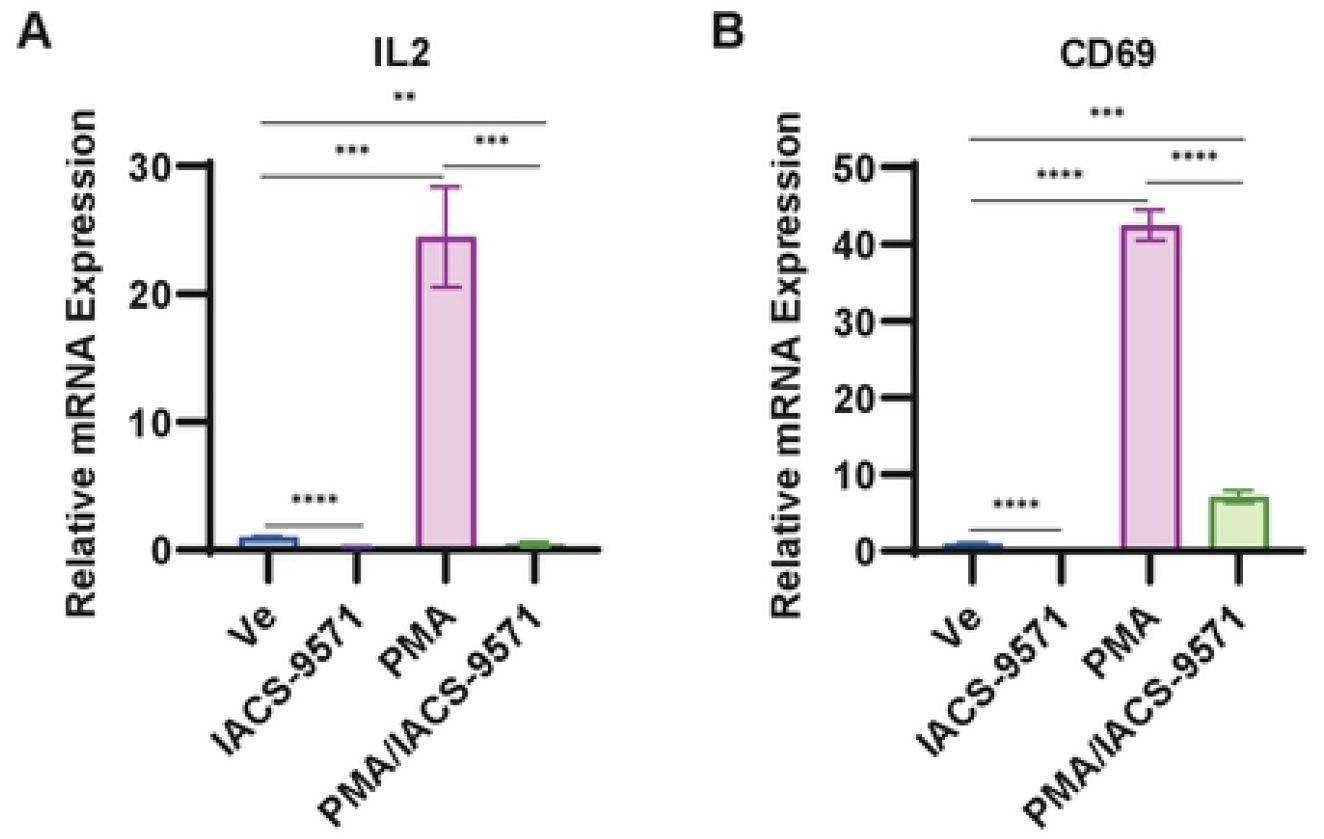
IACS-9571 does not cause activation of Jurkat T cells. **Panel A, B:** Intracellular RNA was extracted from Jurkat T cells following 4 hrs treatment with DMSO (Ve), 10 μM IACS-9571, 20 nM PMA, or 10 μM IACS-9571 and 20 nM PMA. RT-PCR was performed using oligos targeting IL2 (Panel A) or CD69 (Panel B) mRNA. Displayed are the results of three RT-PCR measurements with standard deviation shown.

